# The Future is 2D: Spectral-Temporal Fitting of Dynamic Magnetic Resonance Spectroscopy Data Provides Exponential Gains in Precision Over Conventional Approaches

**DOI:** 10.1101/2022.06.23.497318

**Authors:** Assaf Tal

## Abstract

**Purpose:** Many MRS paradigms produce 2D spectral-temporal datasets, including diffusion-weighted, functional, hyperpolarized and enriched (^13^C, ^2^H) experiments. Conventionally, temporal parameters – such as T_2_, T_1_, or diffusion constants – are assessed by first fitting each spectrum independently, and subsequently fitting a temporal model (1D fitting). We investigated whether simultaneously fitting the entire dataset using a single spectral-temporal model (2D fitting) would improve the precision of the relevant temporal parameter.

**Methods:** We derived a Cramer Rao Lower Bound for the temporal parameters for both 1D and 2D approaches, for two experiments: A multi-echo (MTE) experiment, designed to estimate metabolite T_2_s; And a functional (fMRS) experiment, designed to estimate fractional change (*δ*) in metabolite concentrations. We investigated the dependence of the relative standard deviation of T_2_ in MTE and *δ* in fMRS.

**Results:** When peaks were spectrally distant, 2D fitting improved precision by approximately 20% relative to 1D fitting, regardless of the experiment and other parameter values. These gains increased exponentially as peaks drew closer. Dependence on temporal model parameters was weak to negligible.

**Conclusion:** Our results strongly support a 2D approach to MRS fitting where applicable, and particularly in nuclei such as ^1^H and ^2^H, which exhibit substantial spectral overlap.

## 1. Introduction

Dynamic Magnetic Resonance Spectroscopy refers to an experiment by which a series of one dimensional spectra are acquired sequentially, often while varying a sequence parameter or administering an external time-dependent stimulus or manipulation. This encompasses a wide range of experimental designs, including the observation of the dynamic incorporation of ^13^C or ^2^H-labeled metabolites following the injection of a labeled compound (1–7); Functional MRS, designed to detect endogenous metabolic changes in glutamate, GABA and lactate in response to an external visual, motor or cognitive manipulation (8–20); Multiparametric magnetic resonance spectroscopic experiments, aimed at simultaneously and efficiently quantifying multiple spin parameters (21–26); Diffusion MRS, whereby the diffusion-weighting gradients are varied to quantify the diffusion coefficient of different metabolites (27–32); and even simple relaxometry, where, e.g., T_2_ might be measured by measuring spectral data at different echo times (MTE) (33–41).

The analysis of the two-dimensional spectral-temporal data sets produced by dynamic MRS experiments is conventionally done piece-wise in two stages. First, each spectrum is fit using a linear combination of basis functions (42–48), to extract the temporal dependence of each metabolite’s amplitudes. Then, the time-series for each metabolite’s amplitude is fit to the dynamic model which describes the temporal behavior to extract the relevant temporal constants, such as the transverse or longitudinal relaxation time, diffusion coefficient or metabolite kinetics, depending on the experiment in question. We will refer to this approach as piece-wise, or one-dimensional (1D). Recently, it has been suggested that multiple spectra comprising a dynamic data set should be analyzed and fitted in tandem rather than sequentially, using a model which combines the spectral and temporal degrees of freedom (42). Such an approach utilizes the temporal correlations inherent in the data to benefit the spectral estimations of metabolite amplitudes, and - in principle - should provide more precise and accurate estimates of the temporal constants. We will refer to such approaches as dynamic, or two-dimensional (2D). The two approaches are contrasted schematically in Fig. 1.

**Fig. 1.**
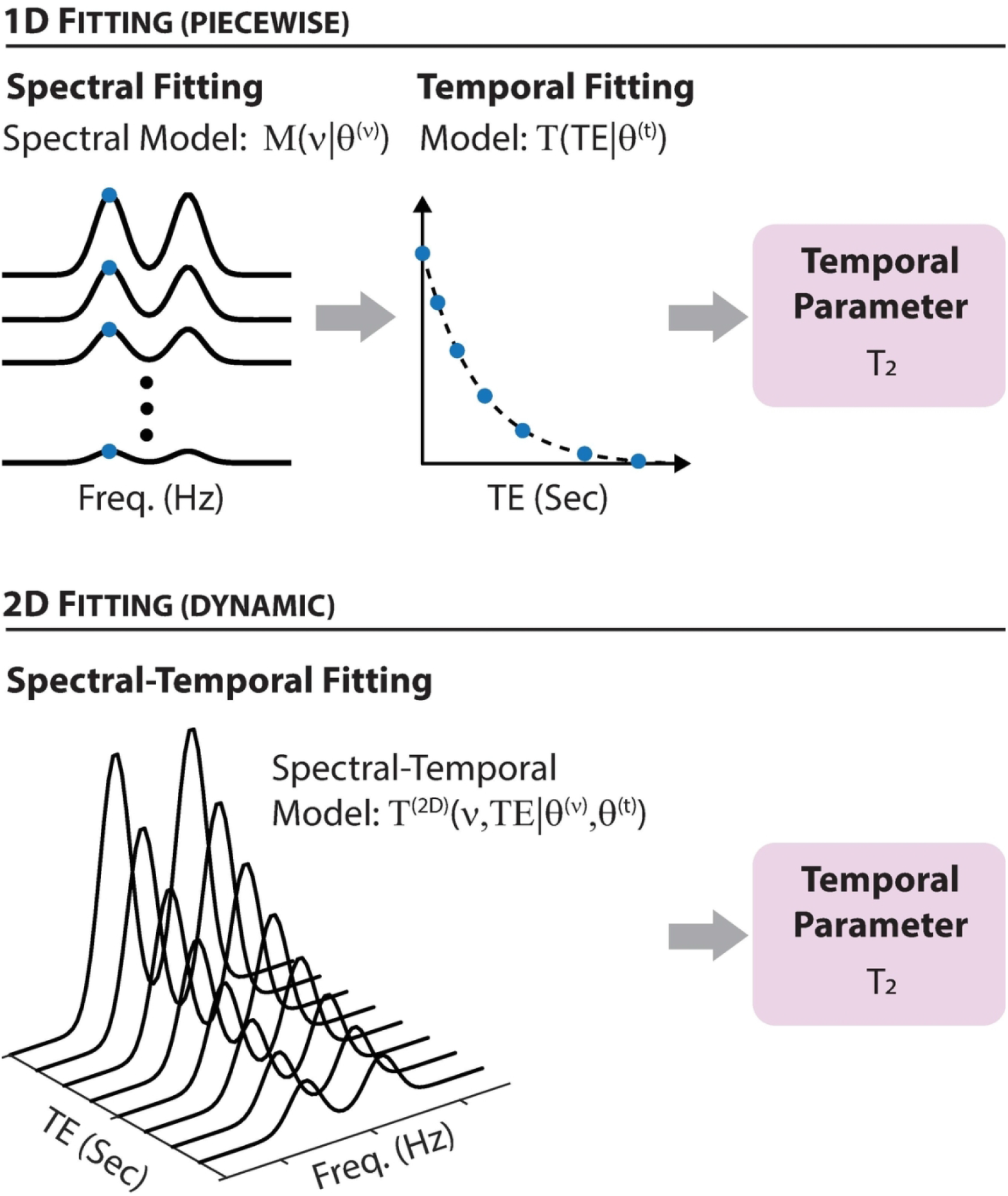
A schematic overview of 1D (piecewise) vs 2D (dynamic) fitting schemes, for a fictitious multi-echo MRS experiment designed to estimate the transverse relaxation time. **Top**: In 1D fitting, data is initially fit to a spectral model to extract the relevant spectral parameters, *θ*^(*v*)^, such as the area (or amplitude) of each spectral peak or metabolite basis function. These are then fit to the relevant temporal model, to extract the relevant temporal parameters, *θ*^(*t*)^ - which, for a multi-echo experiment, include the transverse relaxation time T_2_. **Bottom**: In 2D fitting, the entire spectral-temporal dataset is fit simultaneously to a spectral-temporal model. Such fitting simultaneously furnishes all spectral and temporal parameters - e.g., T_2_ for a multi-echo experiment.. 112×134mm (300 × 300 DPI)

In the current work, we set out to investigate the theoretical gains in precision offered by 2D fitting of dynamic MRS data, relative to the more conventional 1D approach. Rather than quantifying the exact improvements, which would invariably depend on the specific details of the temporal and spectral models, we instead asked ourselves two questions: First, is 2D fitting indeed uniformly superior to 1D fitting? And, if so, which specific spectral or temporal features - or combination thereof - yielded the most substantial gains? To answer these, we assumed a simple spectral model, consisting of two Gaussian peaks, and investigated two temporal models: A multi-echo (MTE) relaxometry experiment designed to estimate T_2_ (which, formally, is equivalent to a diffusion weighted-experiment designed to estimate the apparent diffusion coefficient); And a functional MRS (fMRS) experiment in which one of the peaks changes in response to an external stimulus, while the other remains unchanged. For each model (MTE, fMRS) and each approach (1D, 2D), we calculated the Cramer Rao Lower Bound (49,50), a theoretical estimate on the variance of the relevant dynamical parameter – T_2_ (MTE), and fractional metabolite change (fMRS) - and explored the relative gain in precision offered by 2D fitting.

## Materials and Methods

### Models

Our spectral model ℳ(*v*│*θ*^(*v*)^) consisted of the sum of two Gaussian lineshapes, each with a respective amplitude (A), center (*µ*) and linewidth (Δ):

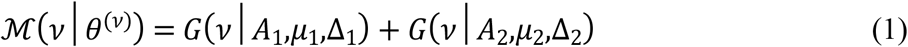

with *θ*^(*v*)^ = (*A*_1_,*µ*_1_,Δ_1_,*A*_2_,*µ*_2_,Δ_2_), and

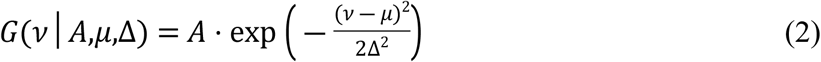

In this notation, *v* is the independent frequency variable, while A, *µ* and Δ are the model parameters.

We considered two temporal models: First, a multi-echo (MTE) experiment, in which the signal decays exponentially with a time constant T_2_ as a function of the echo time (TE):

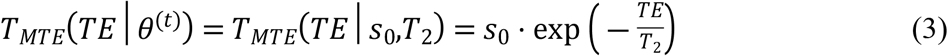

With *θ*^(*t*)^ = (*s*_0_,*T*_2_) the vector of temporal constants. We note that an MTE experiment is formally equivalent to a simple diffusion weighted experiment in which peak amplitudes decay exponentially with a time constant given by the apparent diffusion coefficient D as a function of the b-value, which is altered by changing the diffusion encoding gradient amplitudes (so *θ*^(*t*)^ = (*s*_0_,*D*)). The full 2D dynamic model for each peak consisted of the outer product of the spectral and temporal models, with one minor modification: The amplitude parameter *s*_0_ in the temporal model was set to unity, given that the spectral amplitudes *A*_1_, *A*_2_ can be used to adjust the overall amplitude of each peak:

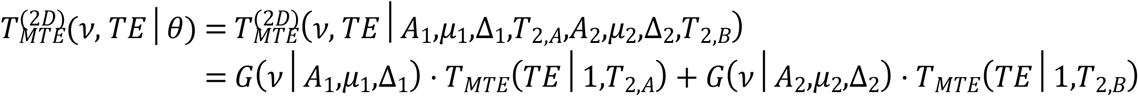

The second temporal model is that of an fMRS experiment using a single-condition block design. Such a design is described the convolution of a boxcar function H(t) of unit amplitude between some initial (*t*_*i*_) and final (*t*_*f*_) times - representing the external stimulus, such as a visual checkerboard pattern - and a point spread function (PSF), which we have taken to be Gaussian with a width 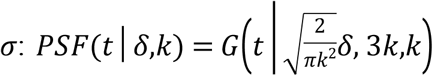. This ensures that the fractional change in the metabolite’s amplitude will be given by *δ*. The shift 3*k* ensures causality, effectively zeroing out the temporal dynamics prior to the administration of the stimulus. Thus:

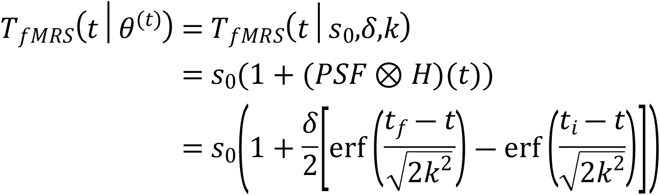

while the full 2D spectral-temporal model is:

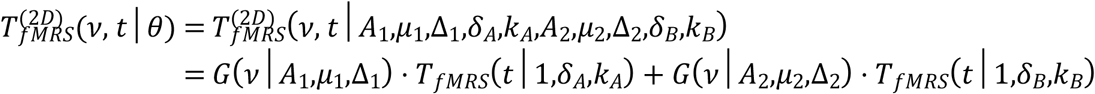

This temporal model can also be used to describe the incorporation of a labeled nucleus – such as ^2^H or ^13^C – in the observable peaks as it undergoes different metabolic cycles within the tissue.

### Cramer Rao Lower Bounds (CRLBs)

For all models we assume the acquired data contains additive, normally distributed noise with zero mean. Under this simplification, the Fisher information matrix for a model with *η* = (*η*_1_(*θ*), *η*_2_(*θ*),…,*η*_*N*_(*θ*)) measurements which depend on M parameters *θ* = (*θ*_1_,*θ*_2_,…,*θ*_*M*_) has matrix elements given by:

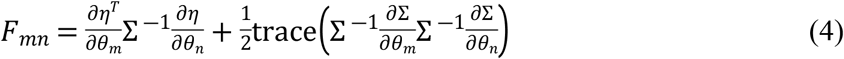

Where Σ is the *N* × *N* covariance matrix of the N measurements.

For the full 2D models, the covariance matrix is diagonal and proportional to the identity, 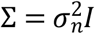, with the noise’s variance being 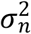. The means are all independent, drawn from the same distribution, and given by the relevant model expression; e.g., for 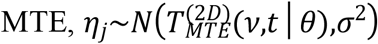. This implies the derivatives of the covariance matrix are all zero and the second term in Eq. (4) can be omitted. The *M* × *M* CRLB matrix is derived by inverting the *M* × *M* Fisher Information matrix, and equals to covariance matrix of the parameters in *θ*. The square root of its diagonal provides the lower bound on the standard deviation of each parameter in *θ*, including the relevant temporal constant (e.g., T_2,A_ and T_2,B_ for MTE). Note that if *N*^(*v*)^ points are sampled along the spectral dimension and *N*^(*t*)^ spectra are acquired along the temporal dimension, then the number of independent measurements is *N* = *N*^(*v*)^*N*^(*t*)^.

For piece-wise 1D fitting approaches, the analysis is slightly more involved: First, the 1D spectral model (Eq. 1) is fitted, and then the amplitude of each peak is fit to the corresponding 1D temporal model to extract the relevant temporal constant. Consequently, Eq. (4) is first computed and inverted to calculate the 6 × 6 covariance matrix of the six spectral parameters in *θ*^(*v*)^ (*σ*_*n*_ is assumed the same as for the full 2D model). The 2 × 2 covariance sub-matrix for the two amplitudes *A*_1_, *A*_2_ is then extracted for each 1D spectrum, and *N*^(*t*)^ such covariance matrices are then chained together along the diagonal to produce a 2*N*^(*t*)^ × 2*N*^(*t*)^ covariance matrix Σ for the temporal model; the block diagonal form reflects the statistical dependence of the two spectral amplitudes, which becomes non-negligible once the peaks overlap. The expression for the mean, *η*_*k*_(*θ*^(*t*)^), is given by the corresponding expression for the 1D temporal model (e.g. Eq. 3 for MTE). Calculating and inverting Eq. 4 produces the covariance matrix of the temporal parameters, and the square root of its diagonal yields the standard deviations of each of the temporal constants.

### Spectral Parameters

Because spectral lines tend to have very similar linewidths, we set the widths of our Gaussian lines to be identical throughout: Δ_1_ = Δ_2_ ≡ Δ = 2.355 · FWHM, where FWHM is the full-width at half maximum of the Gaussian function. Typical spectral lineshapes are approximately 5-10 Hz, and we set FWHM = 8 Hz. For all simulations we defined a distance parameter,

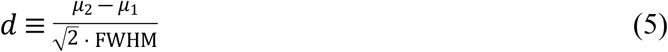

which quantified the closeness of the two Gaussian peaks. *d* ≫ 1 implies little to no overlap, while *d*∼1 and lower signifies significant overlap.

Because the CRLB matrices are always proportional to 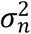, the variance of the noise in the spectra, and because we were only interested in relative standard deviations between the two approaches, the absolute value of *σ*_*n*_ was irrelevant and set to unity. The spectral range was set to [ -32, 32] Hz and sampled at a spectral resolution of 1 Hz, equivalent to a typical FID acquisition time of 1 second, with *N*^(*v*)^ = 64.

### Temporal Parameters: MTE

For MTE, we investigated the relative standard deviation of the two decay constants *T*_2,*A*_ and *T*_2,*B*_ between the two fitting approaches as a function of *d*. First, we considered three cases of equal *T*_2_s: *T*_2,*A*_ = *T*_2,*B*_ = 50, 100 and 200 ms. We also calculated the effect of unequal *T*_2_s, by fixing *T*_2,*A*_ = 100 *ms* and varying *T*_2,*B*_ between 30 and 300 ms, for two values of d: d=0.7 and d=1.4. For all cases, the following echo times were sampled: *TE* = 0,50,100,…,350 ms; This resulted in a 2D dataset with *N*^(*v*)^ × *N*^(*t*)^ = 64 × 8 = 512 data points.

### Temporal Parameters: fMRS

For fMRS, we investigated the relative standard deviation of the fractional change during activation, *δ*, between the two fitting approaches. We assumed a 2-minute experiment with TR=2 seconds and a 60 second boxcar stimulus administered between 30 and 90 seconds. Only one of the peaks was assumed to change, while the other remained static (*δ*_*B*_ ≠ 0, *δ*_*A*_ = 0). We plotted the standard deviation of the fractional change *δ* for each of the two peaks as a function of d for the both fast (k=2 sec) and slow (k=15 sec) metabolite dynamics, for *δ*_*B*_ = 0.05, 0.1 and 0.2. We then explored the effect of the activation timescale, k,, for two spectral peak distances, *d* = 0.8, 1.5, assuming *δ*_*B*_ = 0.1. For each case, the resulting 2D dataset consisted of *N*^(*v*)^ × *N*^(*t*)^ = 64 × 60 = 3840 data points.

## Results

### MTE

The relative gains in precision of T_2_ offered by 2D fitting are summarized in Fig. 3. Fig. 3a shows that the relative SD is independent of T_2_, as long as *T*_2,*B*_ = *T*_2,*A*_. When the peaks have little to no spectral overlap (*d*∼1 and above), this relative gain is 20%; This increases exponentially as the peaks begin to overlap (*d*→0), indicating that 2D fitting could be a powerful tool in resolving the overlap of metabolites in crowded spectra. Fig. 3b shows that some variation (± 20%) in the gain is to be expected when *T*_2,*B*_ ≠ *T*_2,*A*_ and the peaks overlap spectrally (d=0.7), with the optimal performance being observed for *T*_2,*B*_ = *T*_2,*A*_; No variation is observed when the peaks do not overlap (d=1.5). Finally, Fig. 3c – plotted for quality assurance – confirms that the spectral peak amplitudes have no effect on the relative performance, since the CRLB matrix is proportional to the SNR for both 1D and 2D fittings, and this dependence cancels out once the 1D/2D ratio is formed.

**Fig. 2.**
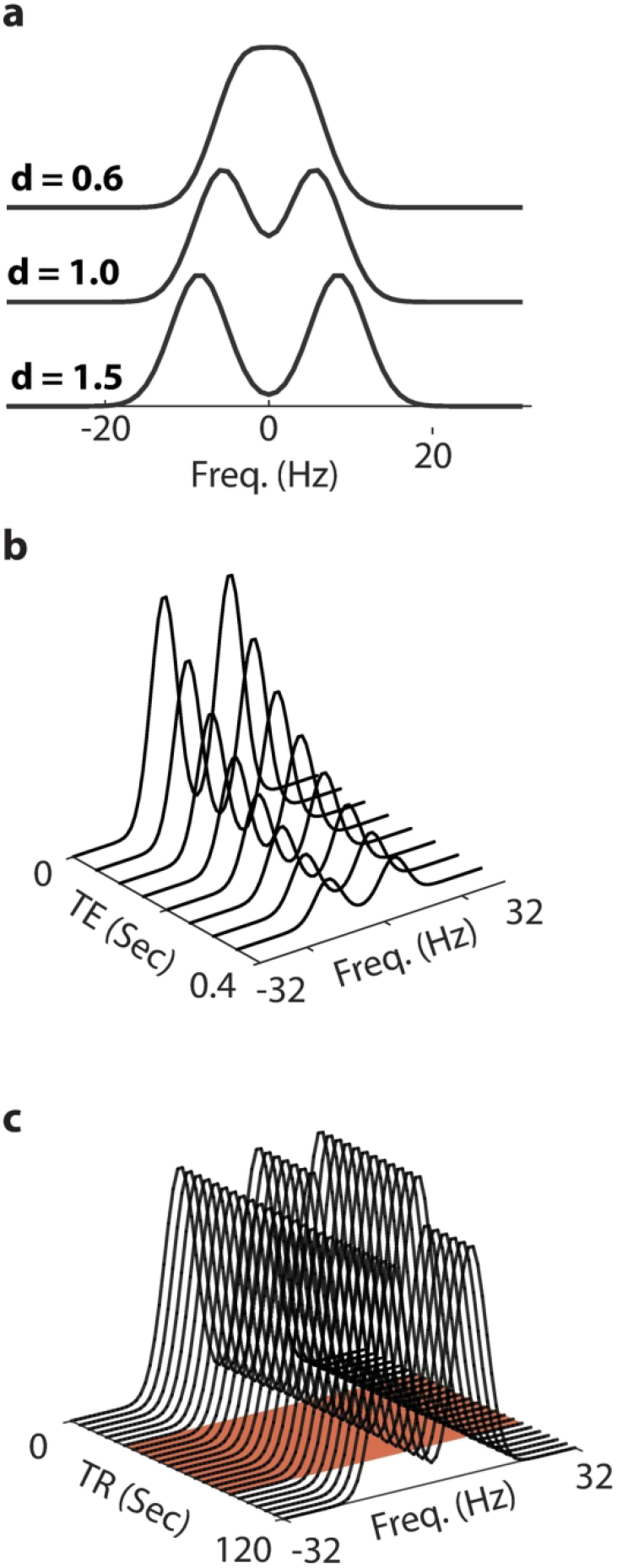
An illustration of the spectral and temporal models under investigation, without any added noise. (a) The spectral model, which consists of two Gaussians, for several values of the distance parameter d (Eq. 5). Additional parameters: unit amplitudes, FWHM of 8 Hz, SW=64 Hz, 64 sampled points. (b) MTE temporal-spectral data, with: Equal amplitudes, FWHM=8 Hz, d=1.5, *T*_2,*A*_ = *T*_2,*B*_ = 200 ms, *TE* = 0,0.05,…0.35 Sec. (c) fMRS temporal-spectral data, with: FWHM=8 Hz, d=1.5, equal amplitudes, *δ*_*A*_ = 0, *δ*_*B*_=0.2, *k*_*A*_ = *k*_*B*_ = 2 seconds, TR=2 seconds. The red-shaded patch indicates the presence of the external stimulus between 30 and 90 seconds. 79×205mm (300 × 300 DPI)

**Fig 3.**
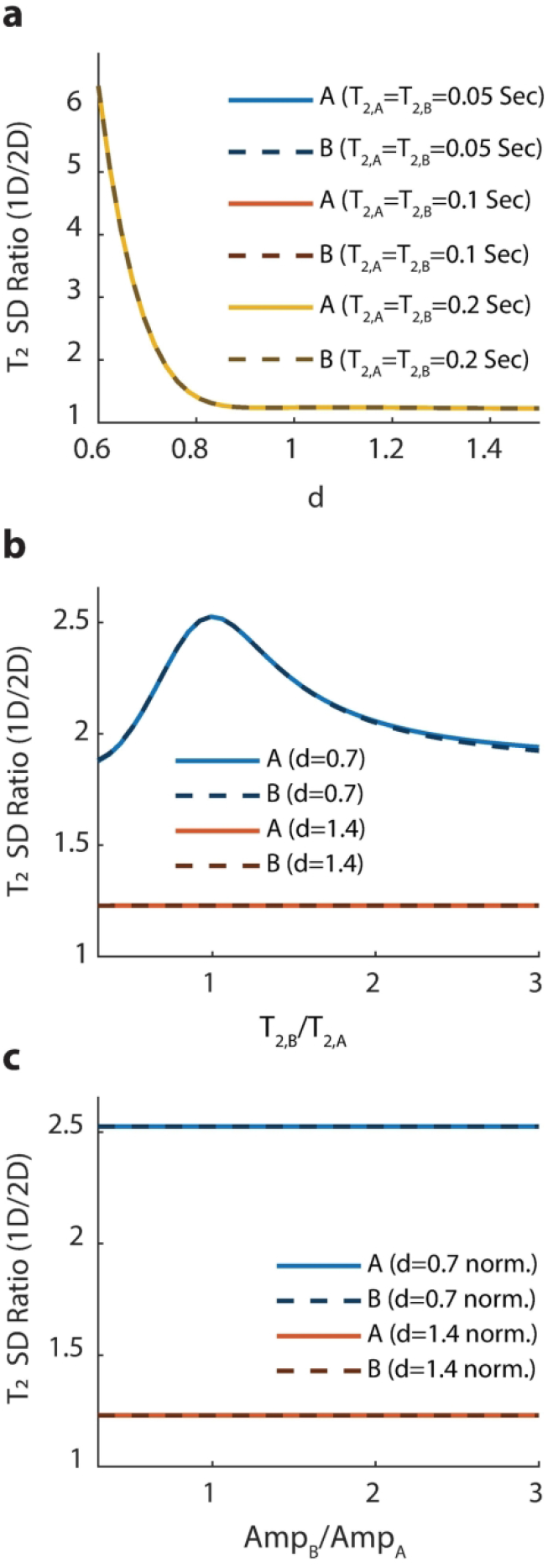
Results: Relative (1D/2D) standard deviation (SD) of T_2_ in an MTE experiment. Solid lines are for peak A while dashed lines are for peak B. Unless otherwise stated, *TE* = 0,0.05,…,0.35 Sec., SW=64 Hz, *N*^(*v*)^ = 64 spectral points, FWHM=8 Hz, and amplitudes are kept equal. (a) Relative SD as a function of the distance d between the spectral peaks, for several values of T_2_, which was kept equal between the two peaks. All curves match almost perfectly, indicating that the relative performance is independent of the absolute value of T_2_ so long as *T*_2,*A*_ = *T*_2,*B*_. This also indicates performance of 2D fitting becomes exponentially better as the peaks begin to spectrally-overlap. (b) Relative SD for two distances (d=0.7, 1.4), while varying the ratio *T*_2,*B*_/*T*_2,*A*_ (fixing *T*_2,*A*_ = 0.1 seconds). When the peaks are far apart, the relative performance is independent of the absolute value of T_2_ (in accordance with (a)). When overlap occurs, 2D fitting provides the best relative performance when the relaxation times are equal. (c) The relative SD a function of the ratio of amplitudes of the two peaks (fixing A_1_=1), for two different spectral distances (d=0.7, 1.4), keeping the amplitude of the first peak fixed at unity. This “sanity check” confirms that, as expected, the performance is independent of the SNR, since the standard deviation of the noise cancels out once the ratio of the CRLBs (1D/2D) is taken. For all plots, as *d* ≫ 1, the relative performance of 2D fitting remains fixed at approximately 1.2 (see Discussion for an explanation of this). 75×217mm (300 × 300 DPI)

### fMRS

The relative gains in precision of *δ*, the fractional change in metabolite concentration, offered by 2D fitting are summarized in Fig. 4. The overall behavior of the relative SD is highly similar between fMRS and MTE: When the peaks do not overlap, the relative gain in precision for 2D fitting is approximately 20%, regardless of all other model parameters; This gain grows exponentially as *d*→0. This is confirmed for both fast (Fig. 4a) and slow (Fig. 4b) temporal dynamics. Fig. 4c shows the ratio of dynamical time constants *k*_*B*_/*k*_*A*_ has a negligible (<5%) effect on the relative SD, whether the peaks overlap spectrally or not.

**Fig 4.**
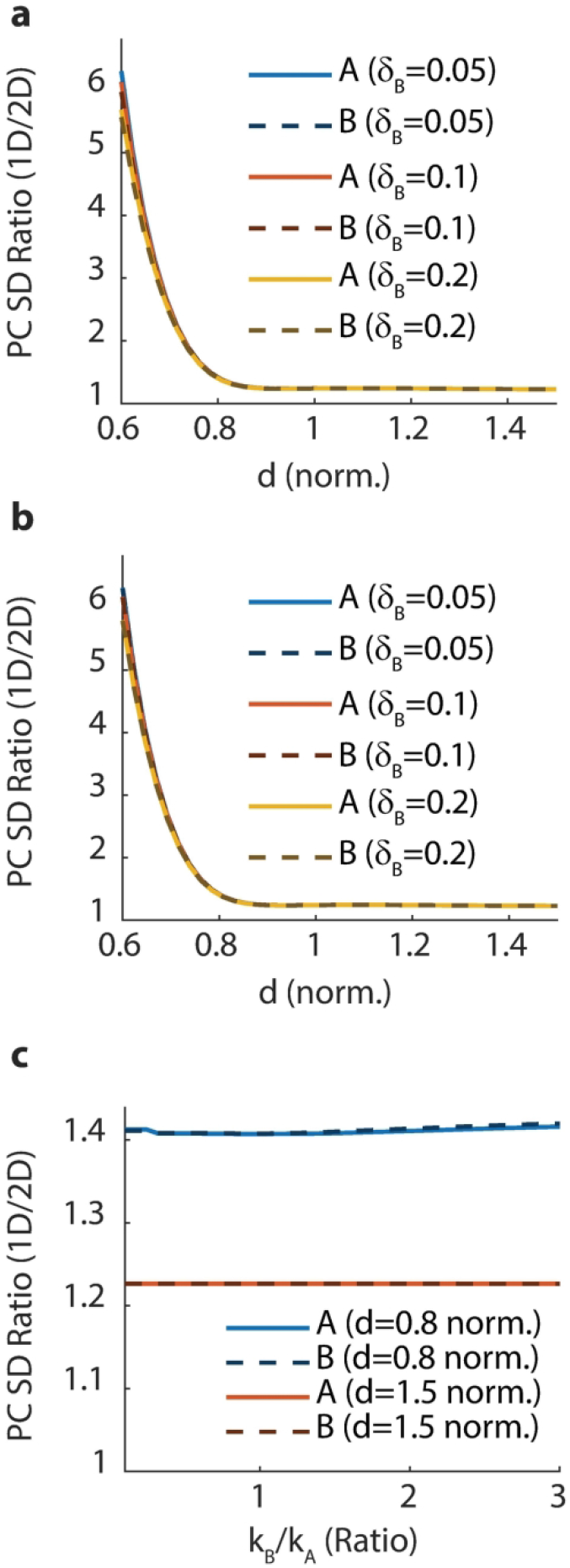
Results: Relative (1D/2D) standard deviation (SD) of the fractional change in metabolite concentrations *δ* in an fMRS experiment. Solid lines are for peak A while dashed lines are for peak B. Unless otherwise stated, TR=2 seconds, TA=2 minutes, and stimulus was administered between 30 and 90 seconds. Spectral parameters were SW=64 Hz, *N*^(*v*)^ = 64 spectral points, FWHM=8 Hz, and equal amplitudes. (a) Relative SD as a function of the spectral distance d between the peaks, for several values of *δ*_*B*_ (keeping *δ*_*A*_ = 0) and fast temporal dynamics (*k*_*A*_ = *k*_*B*_ = 2 seconds). All curves match almost perfectly, indicating that the relative performance is independent of the fractional change. As with the MTE model, this indicates performance of 2D fitting becomes exponentially better as the peaks begin to overlap spectrally. (b) Same as (a), but for slow temporal dynamics (*k*_*A*_ = *k*_*B*_ = 15 seconds). This indicates the temporal dynamics has virtually no effect on the ability to quantify metabolite changes when *k*_*A*_ = *k*_*B*_. (c) Relative SD as a function of the ratio *k*_*B*_/*k*_*A*_, keeping *k*_*A*_ fixed at 5 seconds and varying *k*_*B*_, for two values of the spectral distance (d=0.8, 1.5). This indicates that the ratio of temporal constants has non-zero but negligible (<5%) effect on quantification precision, even when the peaks overlap. Much like for the MTE plots (Fig. 3), the relative gain in precision for 2D fitting remains fixed at approximately 1.2 as *d* increases. 79×221mm (300 × 300 DPI)

## Discussion

In the Introduction we’ve laid out two questions which we are now poised to answer for the two temporal models considered herein (MTE, fMRS): Is 2D fitting always superior to 1D fitting? And when does it achieve its biggest gains? First, our results confirm that 2D fitting is indeed uniformly superior to 1D fitting. For all parameter combinations considered and for all models, the ratio of standard deviations between 2D and 1D approaches exceed ≈ 1.2. The lower bound was attained for both models once the peaks did not overlap spectrally (*d* ≫ 1), regardless of the value of all other parameters. This can be explained by noting that all 2D models fit one parameter fewer than their 1D counterparts: Namely, the amplitude of the temporal model. 1D approaches initially fit the spectral peak amplitudes, and then reintroduce an amplitude parameter in the temporal model (e.g. s_0_ in Eq. 3). It is this redundancy which is avoided by 2D approaches, in which all scaling is done only once in the combined 2D model.

The precision offered by 2D fitting improved exponentially as the spectral overlap between the peaks increased (*d*→0). This behavior was observed for both MTE and fMRS models, regardless of other parameter values. This strongly supports the notion that 2D fitting is a powerful tool for handling spectra with significant overlap - as is the case for proton and deuterium MRS, but less so for enriched ^13^C or ^31^P MRS. The effect of all other parameters was much less significant: the absolute value of the fractional change (*δ*) in an fMRS experiment, or the *T*_2_ relaxation time in an MTE experiment, has little effect on the relative quantification precision. In particular, as long as *T*_2,*A*_ = *T*_2,*B*_ in an MTE experiment, its absolute value had virtually no effect on precision, and different *T*_2,*B*_/*T*_2,*A*_ ratios (from 0.3 to 3.0) led to only a ± 20% deviation in relative precision.

To put the relative gains of 2D fitting in perspective, we note that an N-fold increase in precision is equivalent to an N-fold increase in SNR. Such gains can reduce the sample size N_s_ required to observe an effect with a given statistical significance *α* and power (1 - *β*) (51):

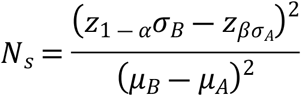

Here *σ*_*B*_, *µ*_*B*_ are the SD and mean of the distribution of a quantity (e.g. percent change, or *T*_2_) in one population (e.g. patients), while *σ*_*A*_, *µ*_*A*_ are the SD and mean in the second population (e.g. controls). z_x_ is the inverse of the normal cumulative distribution function. For typical values of *α* = 0.05, *β* = 0.2, one has *z*_1 − *α*_ = 1.645 and *z*_*β*_ = −0.842. Reducing the SD of both populations by a factor two reduces the required sample size by a factor of four – a substantial gain in experimental time, cost and complexity (however, it should be noted that in a real experiment, *σ*_*A*_, *σ*_*B*_ will be determined by both the intra-subject precision - which 2D fitting improcves - and inter-subject variability, on which 2D fitting has no effect). Even for the case of no spectral overlap, where gains of about 20% in precision are to be expected, a commensurate reduction of 1.2^2^ ≈ 1.44 in the required sample size can be had. Similar substantive gains can be had in sensitivity and specificity, indicating 2D fitting can have a major impact on MRS in both the clinic and in biomedical research.

## Conclusions

By calculating the theoretical CRLB for both 1D and 2D approaches, we have shown that 2D fitting uniformly outperforms 1D fitting, with exponential gains when peaks overlap spectrally Our work strongly motivates the transition to spectral-temporal fitting packages for all dynamic MRS datasets.

## References

1. Feyter HM De Behar KL, Corbin ZA, et al. Deuterium metabolic imaging (DMI) for MRI-based 3D mapping of metabolism in vivo. Sci. Adv. 2018;4:eaat7314 doi: 10.1126/sciadv.aat7314.

2. Lu M, Zhu X-H, Zhang Y, Mateescu G, Chen W. Quantitative assessment of brain glucose metabolic rates using in vivo deuterium magnetic resonance spectroscopy. J. Cereb. Blood Flow Metab. 2017;37:3518–3530 doi: 10.1177/0271678X17706444.

3. Kreis F, Wright AJ, Hesse F, Fala M, Hu D, Brindle KM. Measuring Tumor Glycolytic Flux in Vivo by Using Fast Deuterium MRI. Radiology 2020;294:289–296 doi: 10.1148/radiol.2019191242.

4. Lizarbe B, Lei H, Duarte JMN, Lanz B, Cherix A, Gruetter R. Feasibility of in vivo measurement of glucose metabolism in the mouse hypothalamus by 1H-[13C] MRS at 14.1T. Magn. Reson. Med. 2018;80:874–884 doi: 10.1002/mrm.27129.

5. de Graaf RA, Rothman DL, Behar KL. State of the art direct 13C and indirect 1H-[13C] NMR spectroscopy in vivo. A practical guide. NMR Biomed. 2011;24:958–972 doi: 10.1002/nbm.1761.

6. Rothman DL, de Graaf RA, Hyder F, Mason GF, Behar KL, De Feyter HM. In vivo 13C and 1H-13C MRS studies of neuroenergetics and neurotransmitter cycling, applications to neurological and psychiatric disease and brain cancer. NMR Biomed. 2019;32:e4172 doi: 10.1002/nbm.4172.

7. Rothman DL, de Feyter HM, de Graaf RA, Mason GF, Behar KL. 13C MRS studies of neuroenergetics and neurotransmitter cycling in humans. NMR Biomed. 2011;24:943–957 doi: 10.1002/nbm.1772.

8. Mangia S, Simpson IA, Vannucci SJ, Carruthers A. The in vivo neuron-to-astrocyte lactate shuttle in human brain: evidence from modeling of measured lactate levels during visual stimulation. J. Neurochem. 2009;109:55–62 doi: 10.1111/j.1471-4159.2009.06003.x.

9. Stanley JA, Burgess A, Khatib D, et al. Functional dynamics of hippocampal glutamate during associative learning assessed with in vivo 1H functional magnetic resonance spectroscopy. Neuroimage 2017;153:189–197 doi: 10.1016/j.neuroimage.2017.03.051.

10. Kolasinski J, Hinson EL, Divanbeighi Zand AP, Rizov A, Emir UE, Stagg CJ. The dynamics of cortical GABA in human motor learning. J. Physiol. 2019 doi: 10.1113/JP276626.

11. Woodcock EA, Anand C, Khatib D, Diwadkar VA, Stanley JA. Working Memory Modulates Glutamate Levels in the Dorsolateral Prefrontal Cortex during 1H fMRS. Front Psychiatry 2018;9:66 doi: 10.3389/fpsyt.2018.00066.

12. Ligneul C, Fernandes FF, Shemesh N. High temporal resolution functional magnetic resonance spectroscopy in the mouse upon visual stimulation. Neuroimage 2021;234:117973 doi: 10.1016/j.neuroimage.2021.117973.

13. Jelen LA, King S, Mullins PG, Stone JM. Beyond static measures: A review of functional magnetic resonance spectroscopy and its potential to investigate dynamic glutamatergic abnormalities in schizophrenia. J Psychopharmacol 2018;32:497–508 doi: 10.1177/0269881117747579.

14. Mullins PG. Towards a theory of functional magnetic resonance spectroscopy (fMRS): A meta-analysis and discussion of using MRS to measure changes in neurotransmitters in real time. Scand J Psychol 2018;59:91–103 doi: 10.1111/sjop.12411.

15. Koush Y, de Graaf RA, Jiang L, Rothman DL, Hyder F. Functional MRS with J-edited lactate in human motor cortex at 4T. Neuroimage 2019;184:101–108 doi: 10.1016/j.neuroimage.2018.09.008.

16. Schaller B, Xin L, O’Brien K, Magill AW, Gruetter R. Are glutamate and lactate increases ubiquitous to physiological activation? A 1H functional MR spectroscopy study during motor activation in human brain at 7Tesla. Neuroimage 2014;93:138–145 doi: 10.1016/j.neuroimage.2014.02.016.

17. Schaller B, Mekle R, Xin L, Kunz N, Gruetter R. Net increase of lactate and glutamate concentration in activated human visual cortex detected with magnetic resonance spectroscopy at 7 tesla. J Neurosci Res 2013;91:1076–1083 doi: 10.1002/jnr.23194.

18. Volovyk O, Tal A. Increased Glutamate concentrations during prolonged motor activation as measured using functional Magnetic Resonance Spectroscopy at 3T. Neuroimage 2020;223 doi: 10.1016/j.neuroimage.2020.117338.

19. Taylor R, Neufeld RW, Schaefer B, et al. Functional magnetic resonance spectroscopy of glutamate in schizophrenia and major depressive disorder: anterior cingulate activity during a color-word Stroop task. NPJ Schizophr 2015;1:15028 doi: 10.1038/npjschz.2015.28.

20. Bezalel V, Paz R, Tal A. Inhibitory and excitatory mechanisms in the human cingulate-cortex support reinforcement learning: A functional Proton Magnetic Resonance Spectroscopy study. Neuroimage 2019;184:25–35 doi: 10.1016/j.neuroimage.2018.09.016.

21. Wang CY, Liu Y, Huang S, Griswold MA, Seiberlich N, Yu X. 31P magnetic resonance fingerprinting for rapid quantification of creatine kinase reaction rate in vivo. NMR Biomed. 2017;30 doi: 10.1002/nbm.3786.

22. Lim S-I, Widmaier MS, Wenz D, Yun J, Xin L. Time-efficient relaxation measurements by 31P MR fingerprinting in human brain at 7T. bioRxiv 2022 doi: 10.1101/2022.05.23.493067.

23. Kirov II, Tal A. Potential clinical impact of multiparametric quantitative MR spectroscopy in neurological disorders: A review and analysis. Magn. Reson. Med. 2020;83:22–44 doi: 10.1002/mrm.27912.

24. An L, Li S, Shen J. Simultaneous determination of metabolite concentrations, T1 and T2 relaxation times. Magn Reson Med 2017;78:2072–2081 doi: 10.1002/mrm.26612.

25. Kulpanovich A, Tal A. What is the optimal schedule for multiparametric MRS? A magnetic resonance fingerprinting perspective. NMR Biomed. 2021 doi: 10.1002/nbm.4196.

26. Kulpanovich A, Tal A. The application of magnetic resonance fingerprinting to single voxel proton spectroscopy. NMR Biomed. 2018;31:e4001 doi: 10.1002/nbm.4001.

27. Palombo M, Ligneul C, Najac C, et al. New paradigm to assess brain cell morphology by diffusion-weighted MR spectroscopy in vivo. Proc. Natl. Acad. Sci. 2016;113:6671–6676 doi: 10.1073/pnas.1504327113.

28. Palombo M, Shemesh N, Ronen I, Valette J. Insights into brain microstructure from in vivo DW-MRS. Neuroimage 2018;182:97–116 doi: 10.1016/j.neuroimage.2017.11.028.

29. Branzoli F, Ercan E, Valabrègue R, et al. Differentiating between axonal damage and demyelination in healthy aging by combining diffusion-tensor imaging and diffusion-weighted spectroscopy in the human corpus callosum at 7 T. Neurobiol. Aging 2016;47:210–217 doi: 10.1016/j.neurobiolaging.2016.07.022.

30. Shemesh N, Rosenberg JT, Dumez J-N, Grant SC, Frydman L. Distinguishing neuronal from astrocytic subcellular microstructures using in vivo Double Diffusion Encoded 1H MRS at 21.1 T Motta A, editor. PLoS One 2017;12:e0185232 doi: 10.1371/journal.pone.0185232.

31. Wood ET, Ercan AE, Branzoli F, et al. Reproducibility and optimization of in vivo human diffusion-weighted MRS of the corpus callosum at 3T and 7T. NMR Biomed. 2015;28:976–987 doi: 10.1002/nbm.3340.

32. Genovese G, Marjańska M, Auerbach EJ, et al. In vivo diffusion-weighted MRS using semi-LASER in the human brain at 3 T: Methodological aspects and clinical feasibility. NMR Biomed. 2021;34:e4206 doi: 10.1002/nbm.4206.

33. Lopez-Kolkovsky AL, Mériaux S, Boumezbeur F. Metabolite and macromolecule T1 and T2 relaxation times in the rat brain in vivo at 17.2T. Magn. Reson. Med. 2016;75:503–514 doi: 10.1002/mrm.25602.

34. Murali-Manohar S, Borbath T, Wright AM, Soher B, Mekle R, Henning A. T2 relaxation times of macromolecules and metabolites in the human brain at 9.4 T. Magn. Reson. Med. 2020;84:542–558 doi: 10.1002/mrm.28174.

35. Kirov II, Liu S, Fleysher R, et al. Brain metabolite proton T2 mapping at 3.0 T in relapsing-remitting multiple sclerosis. Radiology 2010;254:858–866 doi: 10.1148/radiol.09091015.

36. An L, Araneta MF, Victorino M, Shen J. Determination of Brain Metabolite T1 Without Interference From Macromolecule Relaxation. J. Magn. Reson. Imaging 2020;52:1352–1359 doi: 10.1002/jmri.27259.

37. Cudalbu C, Mlynárik V, Xin L, Gruetter R. Comparison of T1 relaxation times of the neurochemical profile in rat brain at 9.4 tesla and 14.1 tesla. Magn. Reson. Med. 2009;62:862–867 doi: 10.1002/mrm.22022.

38. Xin L, Schaller B, Mlynarik V, Lu H, Gruetter R. Proton T1 relaxation times of metabolites in human occipital white and gray matter at 7 T. Magn Reson Med 2013;69:931–936 doi: 10.1002/mrm.24352.

39. Xin L, Gambarota G, Mlynárik V, Gruetter R. Proton T2 relaxation time of J-coupled cerebral metabolites in rat brain at 9.4T. NMR Biomed 2008;21:396–401 doi: 10.1002/nbm.1205.

40. Wright AM, Murali-Manohar S, Borbath T, Avdievich NI, Henning A. Relaxation-corrected macromolecular model enables determination of 1H longitudinal T1-relaxation times and concentrations of human brain metabolites at 9.4T. Magn Reson Med 2022;87:33–49 doi: 10.1002/mrm.28958.

41. Kirov II, Fleysher L, Fleysher R, Patil V, Liu S, Gonen O. Age dependence of regional proton metabolites T2 relaxation times in the human brain at 3 T. Magn. Reson. Med. 2008;60:790–795 doi: 10.1002/mrm.21715.

42. Clarke WT, Stagg CJ, Jbabdi S. FSL-MRS: An end-to-end spectroscopy analysis package. Magn. Reson. Med. 2021;85:2950–2964 doi: 10.1002/mrm.28630.

43. Provencher SW. Automatic quantitation of localized in vivo 1H spectra with LCModel. NMR Biomed. 2001 doi: 10.1002/nbm.698.

44. Oeltzschner G, Zöllner HJ, Hui SCN, et al. Osprey: Open-source processing, reconstruction \& estimation of magnetic resonance spectroscopy data. J. Neurosci. Methods 2020;343:108827 doi: 10.1016/j.jneumeth.2020.108827.

45. Wilson M, Reynolds G, Kauppinen RA, Arvanitis TN, Peet AC. A constrained least-squares approach to the automated quantitation of in vivo 1H magnetic resonance spectroscopy data. Magn. Reson. Med. 2011;65:1–12 doi: 10.1002/mrm.22579.

46. Young K, Soher BJ, Maudsley AA. Automated spectral analysis II: Application of wavelet shrinkage for characterization of non-parameterized signals. Magn. Reson. Med. 1998;40:816–821 doi: 10.1002/mrm.1910400606.

47. Gajdošík M, Landheer K, Swanberg KM, Juchem C. INSPECTOR: free software for magnetic resonance spectroscopy data inspection, processing, simulation and analysis. Sci. Rep. 2021;11:2094 doi: 10.1038/s41598-021-81193-9.

48. Edden RAE, Puts NAJ, Harris AD, Barker PB, Evans CJ. Gannet: A batch-processing tool for the quantitative analysis of gamma-aminobutyric acid–edited MR spectroscopy spectra. J. Magn. Reson. Imaging 2014;40:1445–1452 doi: 10.1002/jmri.24478.

49. Cavassila S, Deval S, Huegen C, van Ormondt D, Graveron-Demilly D. Cramér–Rao bounds: an evaluation tool for quantitation. NMR Biomed. 2001;14:278–283 doi: 10.1002/nbm.701.

50. Zhao B, Haldar JP, Liao C, et al. Optimal experiment design for magnetic resonance fingerprinting: Cramér-rao bound meets spin dynamics. IEEE Trans. Med. Imaging 2019;38:844–861 doi: 10.1109/TMI.2018.2873704.

51. Hoch SE, Kirov II, Tal A. When are metabolic ratios superior to absolute quantification? A statistical analysis. NMR Biomed. 2017;30 doi: 10.1002/nbm.3710.

